# Proposal of a taxonomic nomenclature for the *Bacillus cereus* group which reconciles genomic definitions of bacterial species with clinical and industrial phenotypes

**DOI:** 10.1101/779199

**Authors:** Laura M. Carroll, Martin Wiedmann, Jasna Kovac

## Abstract

The *Bacillus cereus* group comprises numerous closely related species, including bioterrorism agent *B. anthracis,* foodborne pathogen *B. cereus*, and biopescticide *B. thuringiensis*. Differentiating organisms capable of causing illness or death from those used in industry is essential for risk assessment and outbreak preparedness. However, current species definitions facilitate species-phenotype incongruencies, particularly when horizontally acquired genes are responsible for a phenotype. Using all publicly available *B. cereus* group genomes (*n* = 2,231), we show that current genomospecies definitions lead to overlapping species clusters, and that an average nucleotide identity (ANI) genomospecies threshold of ≈92.5 reflects a natural gap in core genome similarity. We propose a taxonomy for the *B. cereus* group which accounts for (i) genomospecies using separable species clusters formed at a threshold of ≈92.5 ANI, and (ii) phenotypes relevant to public health and industry. We anticipate that the proposed nomenclature will remain interpretable to clinicians, without sacrificing genomic species definitions, which can in turn aid in pathogen surveillance, early detection of emerging, high-risk genotypes, and outbreak preparedness. Furthermore, the nomenclatural framework outlined here serves as a model for genomics-based bacterial taxonomy which moves beyond arbitrarily set genomospecies thresholds, while maintaining congruence with phenotypes and historically important species names.

## INTRODUCTION

The *Bacillus cereus* group species complex, also known as *B. cereus sensu lato* (*s.l.*), is a subgroup of closely related species belonging to the genus *Bacillus.* Group members are Gram-positive, spore-forming, and widely distributed throughout the environment.^1^ While closely related from an evolutionary perspective, members of this group vary in their ability to cause disease in humans. Notable members which are considered to be pathogenic include anthrax-causing *B. anthracis*, which has been responsible for outbreaks and bioterrorism attacks around the world,^2–5^ and *B. cereus sensu stricto* (*s.s.*), which is commonly regarded as a foodborne pathogen, but has also been associated with anthrax-like symptoms and other severe infections.^1, 6^ Interspersed among species which are widely regarded as pathogenic are those which have found important roles in agriculture and industry, the most notable of which is biopesticide *B. thuringiensis*.^7, 8^

Prior to 2013, the *B. cereus* group was composed of six closely related species (i.e., *B. anthracis, B. cereus s.s.*, *B. mycoides*, *B. pseudomycoides*, *B. thuringiensis*, and *B. weihenstephanensis*), which had been delineated using various methods, including phenotypic characterization (e.g., production of insecticidal crystal proteins [*B. thuringiensis*], rhizoidal colony morphology [*B. mycoides* and *pseudomycoides*], psychrotolerance and an inability to grow at 43°C [*B. weihenstephanensis*]), 16S rDNA sequencing, and/or DNA-DNA hybridization.^9–11^ However, as whole-genome sequencing (WGS) has become more affordable and accessible, the gold standard for prokaryotic species delineation has migrated to high-throughput, *in silico* average nucleotide identity (ANI)-based methods,^12^ for which two genomes are said to belong to the same genomospecies if they share an ANI value above a threshold. While numerous genomospecies thresholds have been proposed over the years (e.g., 94, proposed by Konstantinidis and Tiedje in 2005,^13^ 95-96 ANI, proposed by Richter and Rosselló-Móra^12^ and later supported by Kim, et al.^14^), a recent survey of pairwise ANI values between 90,000 prokaryotic genomes (including some members of the *B. cereus* group) concluded that a genomospecies threshold of 95 ANI should be adequate for most bacterial species.^15^

ANI-based genomospecies assignment for members of the *B. cereus* group has relied on calculating the pairwise ANI values shared between a genome of interest and the genomes of all published *B. cereus* group type strains, a practice that was used to describe *B. cytotoxicus* and *B. toyonensis* (published in 2013), and *B. wiedmannii* (published in 2016) as novel species.^16–18^ This practice was further employed in 2017, when nine novel *B. cereus* group species (*B. albus, B. luti, B. mobilis, B. nitratireducens, B. pacificus, B. paramycoides, B. paranthracis*, *B. proteolyticus*, and *B. tropicus*) were published,^19^ effectively doubling the number of published *B. cereus* group species from nine to 18.

However, the practice of assigning *B. cereus* group genomes to a genomospecies using *B. cereus* group type strain genomes is problematic due to the fact that (i) type strain genomes do not necessarily (or even likely) represent the medoid of a genomospecies cluster, meaning that it is possible for a genome to share an ANI value greater than the genomospecies threshold with multiple type strain genomes (i.e., a genome could potentially belong to more than one *B. cereus* group genomospecies), and (ii) novel *B. cereus* group species have been published using different genomospecies thresholds (e.g., *B. toyonensis, B. wiedmannii*, and the nine species published in 2017 used ANI genomospecies thresholds of 92, 95, and 96, respectively).^17^ Further confusion arises when the type strains of “novel” species encompass well-established, previously described clinically and industrially relevant *B. cereus* group lineages within their genomospecies thresholds. For example, since the publication of *B. paranthracis* as a novel species in 2017,^19^ the well-researched foodborne pathogen referred to as emetic “*B. cereus*”^1, 20–22^ technically belongs to the *B. paranthracis* genomospecies cluster based on a conventional ANI threshold of 95.^23^ This is problematic, as this taxonomic assignment likely bears little meaning to anyone not well-versed and up-to-date with *B. cereus* group taxonomy, including clinicians. Current definitions of *B. cereus* group species are further proven to be outdated as the amount of publicly available genomic data grows and continues to reveal increasing genomic and phenotypic diversity within the group. Between April 2017^24^ and March 2018, the number of assembled *B. cereus* group genomes available in the National Center for Biotechnology Information’s (NCBI’s) RefSeq^25^ database more than tripled, implying that there are likely unexplored portions of the *B. cereus* group phylogeny.

Genomic and taxonomic semantics aside, phenotypic characteristics used for species assignment (e.g., motility, hemolysis, emetic toxin production) are known to vary within and among species.^9, 10, 26, 27^ This is particularly problematic in cases where the genomic determinants responsible for a clinically or industrially relevant phenotype are plasmid-mediated, such as synthesis of anthrax toxin/capsular proteins,^28–31^ bioinsecticidal crystal proteins,^32–34^ or cereulide (emetic toxin) synthetase proteins.^35, 36^ These traits can be lost or gained, heterogeneous in their presence within a species, or present across multiple species.

Current species definitions do not account for species-phenotype incongruencies, which can lead to potentially high-consequence misclassifications of an isolate’s virulence potential. For example, strains exhibiting phenotypic characteristics associated with “*B. cereus*”, such as motility, can cause anthrax in humans and animals^37–40^, while *B. anthracis* which lack the genes required for anthrax toxin and capsule formation have attenuated virulence.^41^ The problem at hand requires the construction of an ontological framework which is accurate in terms of its adherence to widely accepted genomic and taxonomic definitions of bacterial genomospecies, while still being informative, intuitive, and actionable to those in public health and industry. Differentiating organisms capable of causing illness or death in humans and animals from those which have far-reaching agricultural and industrial applications is essential for a proper assessment of the risk posed by a particular strain. Here, we leverage all publicly available assembled *B. cereus* group genomes (*n* = 2,231) to construct a phylogenomically informed taxonomic framework with the flexibility to account for phenotypes of interest to those in public health and industry.

## METHODS

### Acquisition and initial *in silico* characterization of *Bacillus cereus* group genomes

All genomes in the NCBI RefSeq Assembly database^25^ which were submitted as a published *B. cereus* group species (*B. albus, anthracis, cereus, cytotoxicus, luti, mobilis, mycoides, nitratireducens, pacificus, paramycoides, paranthracis, proteolyticus, pseudomycoides, thuringiensis, toyonensis, tropicus, weihenstephanensis,* or *wiedmannii*) were downloaded (Supplementary Tables S1 and S2), along with the type strain genomes of three proposed effective *B. cereus* group species (i.e., *“B. bingmayongensis”, “B. gaemokensis”,* and *“B. manliponensis”*) (*n* = 2,231, accessed November 19, 2018; Supplementary Tables S1 and S2). QUAST version 4.0^42^ was used to assess the quality of each assembled genome, and BTyper version 2.3.2^24^ was used to detect *B. cereus* group virulence genes in each genome, using default minimum amino acid sequence identity and coverage thresholds (50 and 70%, respectively), which have been shown to correlate with PCR-based detection of virulence genes in *B. cereus* group isolates (Supplementary Table S1).^24, 43^ Prokka version 1.12^44^ was used to annotate each of the 2,231 *B. cereus* group genomes, and the resulting coding sequences (CDS) were used as input for the command-line implementation BtToxin_scanner version 1.0 (BtToxin_scanner2.pl),^45^ which was used to identify insecticidal toxin genes associated with *B. thuringiensis* (Bt toxins) in each genome using the default settings.

### Calculation of pairwise average nucleotide identity values, hierarchical clustering, and identification of medoid genomes

FastANI version 1.0^15^ was used to calculate pairwise average nucleotide identity (ANI) values between each of the 2,231 genomes (4,977,361 total comparisons). To ensure that the breakpoints and shape of the distribution of pairwise ANI calculations were robust to genome ambiguity, all pairwise ANI values were calculated a second time, with ambiguous nucleotides (i.e., those not belonging to the set {*𝐴, 𝐶, 𝐺, 𝑇*}) removed from each genome (Supplementary Figure S1). Robustness was further assessed by removing genomes (i) falling below various N50 thresholds (i.e., ≤ 10 Kbp, 20 Kbp, 50 Kbp, and 100 Kbp), and/or (ii) containing any contigs classified in domains other than Bacteria, phyla other than Firmicutes, and/or genera other than *Bacillus* using Kraken version 2.0.8-beta^46, 47^ and the complete standard Kraken database (accessed August 6, 2019; Supplementary Figure S1). For each data set, a histogram of all pairwise ANI values was plotted in R version 3.6.0,^48^ using the ggplot2 package (Supplementary Figure S1).^49^ For the identification of a final set of medoid genomes at various thresholds (described below), all genomes with an N50 > 20 Kbp in the original set of 2,231 NCBI RefSeq genomes were used in all subsequent steps (*n* = 2,218; Supplementary Table S1 and Supplementary Figure S1).

For each data set, the resulting pairwise ANI values were used to construct a similarity matrix, **𝐒_𝐀𝐍𝐈_**, using R version 3.6.0 and the reshape2 package^50^ as follows, where *n* = 2,218: Let *𝑔*_1_, *𝑔*_2_, … *𝑔_n_* be a set of *𝑛* genomes, denoted by *𝐺* (*𝐺* = {*𝑔*_1_, *𝑔*_2_, … *𝑔_n_*}). Similarity function 𝐴𝑁𝐼(*𝑔_i_*, *𝑔j*) denotes the ANI value shared by query and reference genomes *𝑔_i_* and *𝑔j*, respectively, where

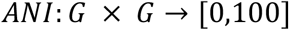

Similarity matrix **𝐒_𝐀𝐍𝐈_** can be defined as

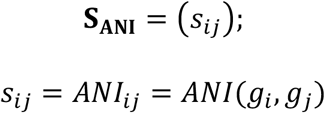

Similarity matrix **𝐒_𝐀𝐍𝐈_** was converted to a dissimilarity matrix, **𝐃_𝐀𝐍𝐈_**, as follows, where 𝐉 denotes an *𝑛* × *𝑛* matrix where each element is equal to 1:

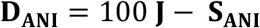

*ANI* as a similarity function is not symmetric (i.e., for all *𝑔_i_*, *𝑔_j_*, 𝐴𝑁𝐼(*𝑔_i_*, *𝑔_j_*) ≠ 𝐴𝑁𝐼(*𝑔_j_*, *𝑔_i_*)), as minor differences between corresponding values in the upper and lower triangles of **𝐃_𝐀𝐍𝐈_** existed: max (*𝑑*(*𝑔_i_*, *𝑔_j_*), *𝑑*(*𝑔_j_*, *𝑔_i_*)) = 0.504, min (*𝑑*(*𝑔_i_*, *𝑔_j_*), *𝑑*(*𝑔_j_*, *𝑔_i_*)) = 0, mean (*𝑑*(*𝑔_i_*, *𝑔_j_*), *𝑑*(*𝑔_j_*, *𝑔_i_*)) = 0.056, and median (*𝑑*(*𝑔_i_*, *𝑔_j_*), *𝑑*(*𝑔_j_*, *𝑔_i_*) = 0.046. As such, **𝐃_𝐀𝐍𝐈_** is not a symmetric matrix (i.e., **𝐃_𝐀𝐍𝐈_** ≠ **𝐃_𝐀𝐍𝐈_^𝐓^**). To coerce **𝐃_𝐀𝐍𝐈_** to a symmetric matrix, 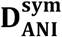, the following transformation was applied:

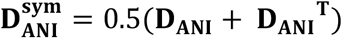

The hclust function in R’s stats package was then used to perform average linkage hierarchical clustering, using 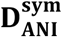 as the dissimilarity structure, and the resulting dendrogram was annotated using the ggplot2, dendextend,^51^ and viridis^52^ packages. Dendrogram clusters formed at various species thresholds (denoted here by *𝑇_d_*, where *𝑇_d_* = {4,5,6,7.5}, corresponding to ANI values of 96, 95, 94, and 92.5, respectively) were obtained by treating genome lineages which coalesced prior to *𝑇_d_* as members of the same cluster (i.e., genomospecies), and those which did not as members of different clusters. Medoid genomes were then identified within each cluster at each threshold, using the pam function in R’s cluster package^53^ and 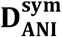 as a dissimilarity structure, where the medoid genome is defined as

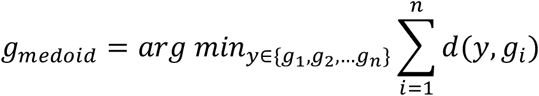

where *𝑑*(*𝑔_i_*, *𝑔_j_*) = 100 − 𝐴𝑁𝐼(*𝑔_i_*, 𝑔*_j_*).

To construct a graph with each of the final set of 2,218 *B. cereus* group genomes represented as nodes and ANI values represented as weighted edges, 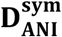 was converted to a symmetric similarity matrix, 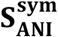, as follows:

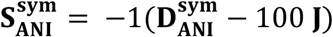

The igraph^54^ package in R version 3.6.0 was used to construct each graph, with 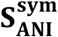 treated as an adjacency matrix, and edges with weights (i.e., ANI values) less than a similarity threshold *𝑇_S_* (i.e., *𝑇_S_* = {92.5,95}) removed.

### Phylogeny construction using single-copy core orthologous clusters identified in 2,231 *B. cereus* group genomes

FASTA files containing amino acid sequences of protein-coding features (.faa files) produced by Prokka version 1.12^44^ were used as input for OrthoFinder version 2.3.3.^55^ Single-copy orthologous clusters present in all 2,231 genomes (i.e., single copy core orthologous clusters) were identified using an iterative approach, in which OrthoFinder was used to identify single-copy orthologous clusters core to *n* of the 2,231 *B. cereus* group genomes, sampled randomly without replacement (where *n* = 30 or *n* = 11 for 74 and 1 [the remainder] iteration[s], respectively). The union of single-copy orthologous clusters present in all *n* genomes in each random sample of *B. cereus* group genomes was then queried again using OrthoFinder, which identified a total of 79 single-copy orthologous clusters core to all 2,231 *B. cereus* group genomes. Nucleotide sequences of each of the 79 single-copy core orthologous clusters present in all 2,231 genomes were aligned using PRANK v.170427.^56^ The resulting alignments were concatenated, and snp-sites version 2.4.0^57^ was used to produce an alignment of variant sites, excluding gaps and ambiguous characters. IQ-TREE version 1.6.10^58^ was used to construct a maximum likelihood phylogeny, using the alignment of core SNPs detected in all 2,231 *B. cereus* group genomes. The GTR+G+ASC nucleotide substitution model implemented in IQ-TREE (i.e., General Time Reversible model^59^ with a Gamma parameter^60^ to allow rate heterogeneity among sites and an ascertainment bias correction^61^ to account for the use of solely variant sites) was used, along with 1,000 replicates of the Ultrafast bootstrap approximation.^62^ Taxa excluded from the final medoid set of genomes (i.e., those with N50 < 20 Kbp) were removed using the drop.tip function in the ape package for R version 3.6.0, and the resulting phylogeny was annotated in R using the following packages: ggplot2, ape,^63^ phytools,^64^ phylobase,^65^ ggtree,^66^ and phangorn.^67^

## RESULTS

### Current species definitions do not reliably differentiate *B. anthracis* from neighboring lineages

The currently employed practice of calculating pairwise ANI values between a genome of interest and the genomes of known *B. cereus* group species type strains (Supplementary Table S2)^17–19, 23^ and using the widely accepted threshold of 95 ANI^15^ as a hard genomospecies cutoff produced non-overlapping, separable genomospecies clusters for *B. albus* (*n* = 11), “*B. bingmayongensis*” (*n* = 1), *B. cytotoxicus* (*n* = 14), “*B. gaemokensis*” (*n* = 1), *B. luti* (*n* = 3), “*B. manliponensis*” (*n* = 1), *B. nitratireducens* (*n* = 70), *B. paramycoides* (*n* = 6), *B. proteolyticus* (*n* = 7), *B. pseudomycoides* (*n* = 111), and *B. toyonensis* (*n* = 230). None of the genomes assigned to these genomospecies clusters shared ≥95 ANI with any genomes assigned to a different genomospecies cluster (Figures 1 and 2A1 and Supplementary Table S3). However, several currently defined type strain-centric genomospecies clusters did not yield well-separated, reliable genomospecies assignments, including genomospecies clusters formed by the type strains of (i) diarrheal foodborne pathogen *B. cereus s.s.* and biopesticide *B. thuringiensis*, and (ii) *B. mycoides* and *B. weihenstephanensis,* as has been well-documented previously (Figures 1 and 2A1 and Supplementary Table S3).^18, 68, 69^ The type strains of newly described *B. mobilis* and *B. wiedmannii* (published in 2017 and 2016, respectively)^18, 19^ were also found to produce ambiguous taxonomic classifications in which a genome could share ≥95 ANI with both species type strains (Figures 1 and 2A1 and Supplementary Table S3). The largest source of ambiguity, however, stemmed from bioterrorism agent *B. anthracis* and its neighboring lineages: the genomospecies cluster formed by the *B. anthracis* reference genome overlapped with those of *B. pacificus*, *B. paranthracis,* and *B. tropicus*, the latter three of which were published as novel species in 2017 (Figures 1 and 2A1 and Supplementary Table S3).^19^ While no genomes were found to share ≥95 ANI with three or more of *B. anthracis*, *B. pacificus, B. paranthracis*, and *B. tropicus* genomospecies clusters, each pair of these four genomospecies clusters was found to overlap (Figures 1 and 2A1 and Supplementary Table S3).

**Figure 1.**
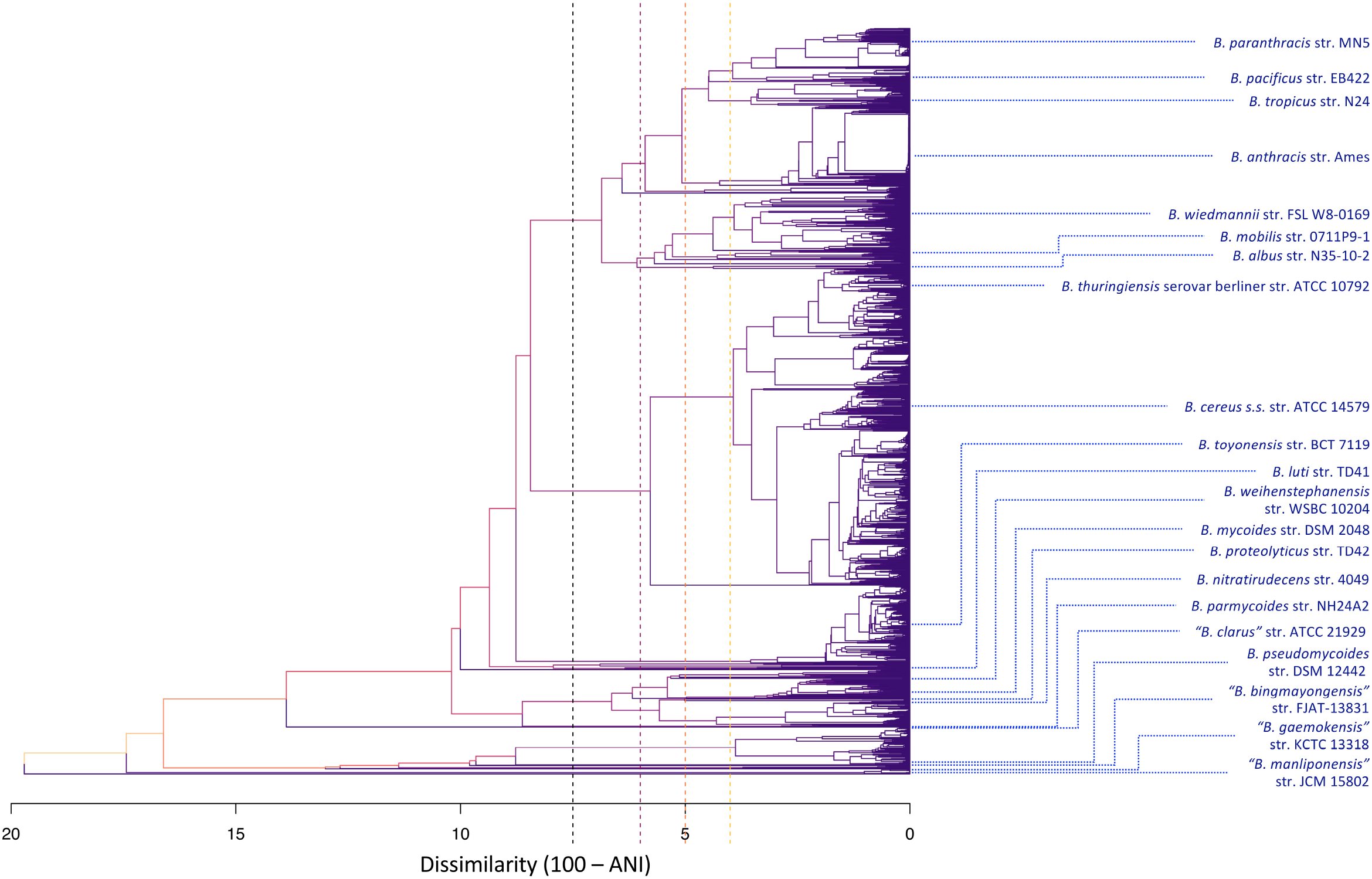
Dendrogram constructed using symmetric pairwise average nucleotide identity (ANI) dissimilarities calculated between 2,218 *B. cereus* group genomes from NCBI’s RefSeq database with N50 > 20 Kbp (i.e., 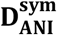 in the “Methods” section) and the average linkage hierarchical clustering method implemented in the hclust function in R. Blue tip labels denote the location of species type strain/reference genomes in the dendrogram, while tree height corresponds to ANI dissimilarity. Branch colors correspond to branch height within the tree. Dashed vertical lines appear at dissimilarities of 7.5, 6, 5, and 4, which correspond to ANI thresholds of 92.5, 94, 95, and 96, respectively (from left to right in order of appearance along the X-axis).

The species overlap problem persisted at a 95 ANI threshold, even when medoid genomes were used to represent genomospecies clusters instead of type strain genomes (Figure 2A2 and Supplementary Table S4). All genomospecies clusters which were non-overlapping when type strains were used for genomospecies assignment (e.g., *B. pseudomycoides, B. toyonensis*) remained non-overlapping, except for *B. proteolyticus*, which was located at the intersection of two genomospecies clusters (Figure 2A2 and Supplementary Table S4). All overlapping genomospecies clusters (i.e., *B. cereus s.s.* and *B. thuringiensis*; *B. mycoides* and *B. weihenstephanensis*; *B. mobilis* and *B. wiedmannii*; *B. anthracis, B. pacificus, B. paranthracis,* and *B. tropicus*) continued to produce ambiguous genomospecies assignments, although with more than 3.6 times fewer total multi-species classifications at ≥ 95 ANI compared to assignment based on species type strains; 405 genomes were assigned to 2 or more medoid-based genomospecies clusters [18.2%], compared to 1,478 genomes assigned to 2 or more type strain genomospecies clusters [66.2%] (Figure 2A).

**Figure 2.**
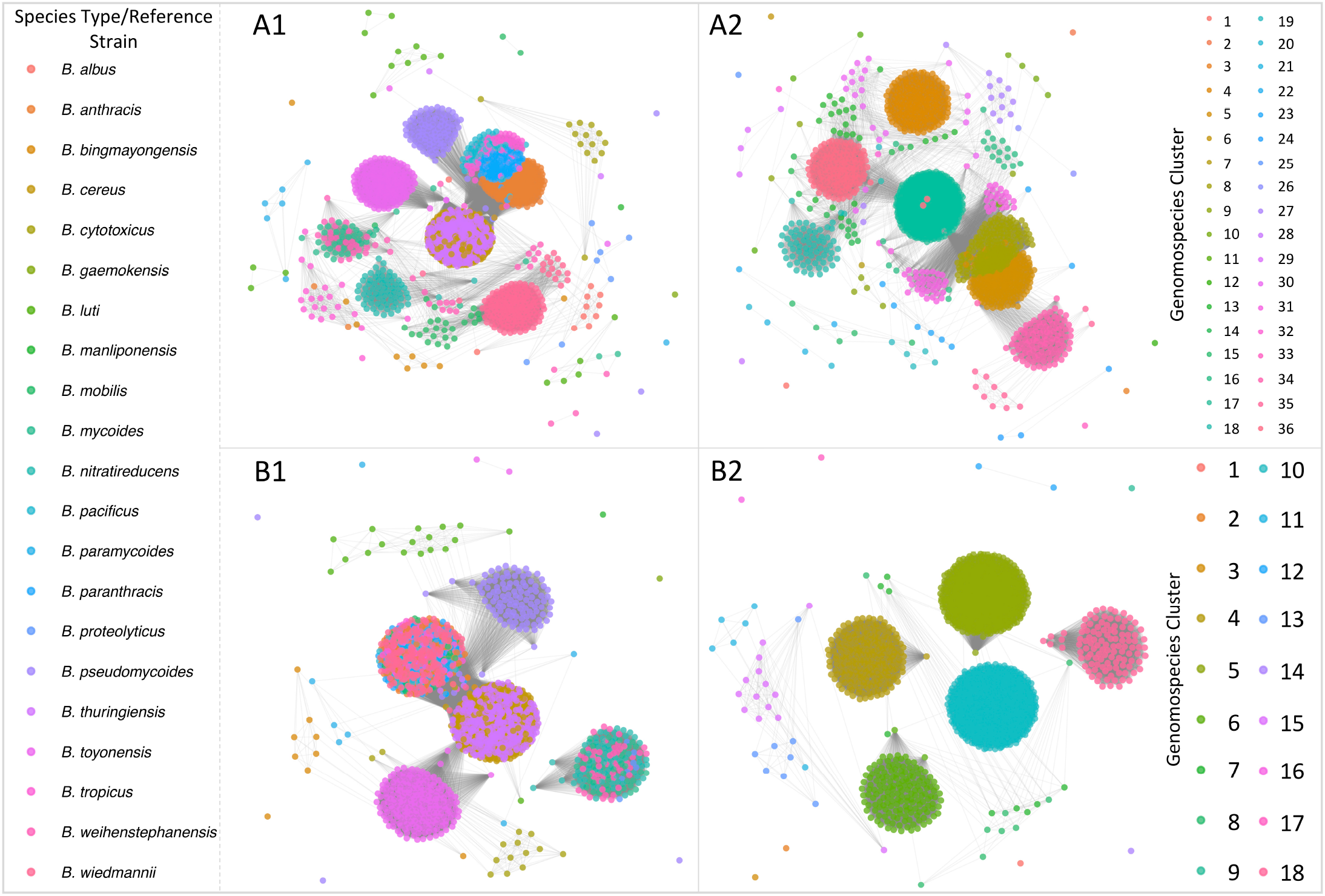
Weighted undirected graphs constructed using symmetric pairwise average nucleotide identity (ANI) values calculated between 2,218 *B. cereus* group genomes from NCBI’s RefSeq database with N50 > 20 Kbp (i.e., 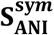 in the “Methods” section). Nodes represent individual genomes, while weighted edges connect each pair of genomes with a mean ANI value (A) ≥ 95, and (B) ≥ 92.5, where edge weight corresponds to the mean ANI value of the pair. Nodes (i.e., genomes) are colored by (1) closest matching type strain genome, or (2) closest matching medoid genome of clusters formed at the respective ANI value. Graphs were constructed using the graphout layout algorithm implemented in R’s igraph package, using 500 iterations and a charge of 0.02.

### Genomic elements responsible for anthrax, emetic, and insecticidal toxin production exhibit heterogeneous presence in multiple *B. cereus* group species using current genomospecies definitions

Additional nomenclatural discrepancies arise when a trait of interest is plasmid-encoded, as these traits are expected to be lost or gained more frequently than those of chromosomal origin. Such is the case of the plasmid-mediated anthrax toxin genes often associated with *B. anthracis* (edema factor-encoding *cya*, lethal factor-encoding *lef* and protective antigen-encoding *pagA*):^70^ 93 of 241 (38.6%) genomes most closely resembling the *B. anthracis* reference genome at ≥95 ANI did not possess anthrax toxin genes *cya*, *lef,* and *pagA* (Figures 3A and 3B and Supplementary Table S5). Notably, isolates which most closely resemble *B. anthracis* by current species definitions (i.e., ≥95 ANI), despite lacking the genes necessary to produce anthrax toxin, do not appear to be particularly uncommon; such strains have been isolated from a diverse array of environments (e.g., soil, animal feed, milk, spices, egg whites, baby wipes), from six continents, as well as the International Space Station (Supplementary Table S5). The classification of these isolates as *B. anthracis* could lead to incorrect assumptions about the anthrax-causing capability of strains belonging to these lineages.

**Figure 3.**
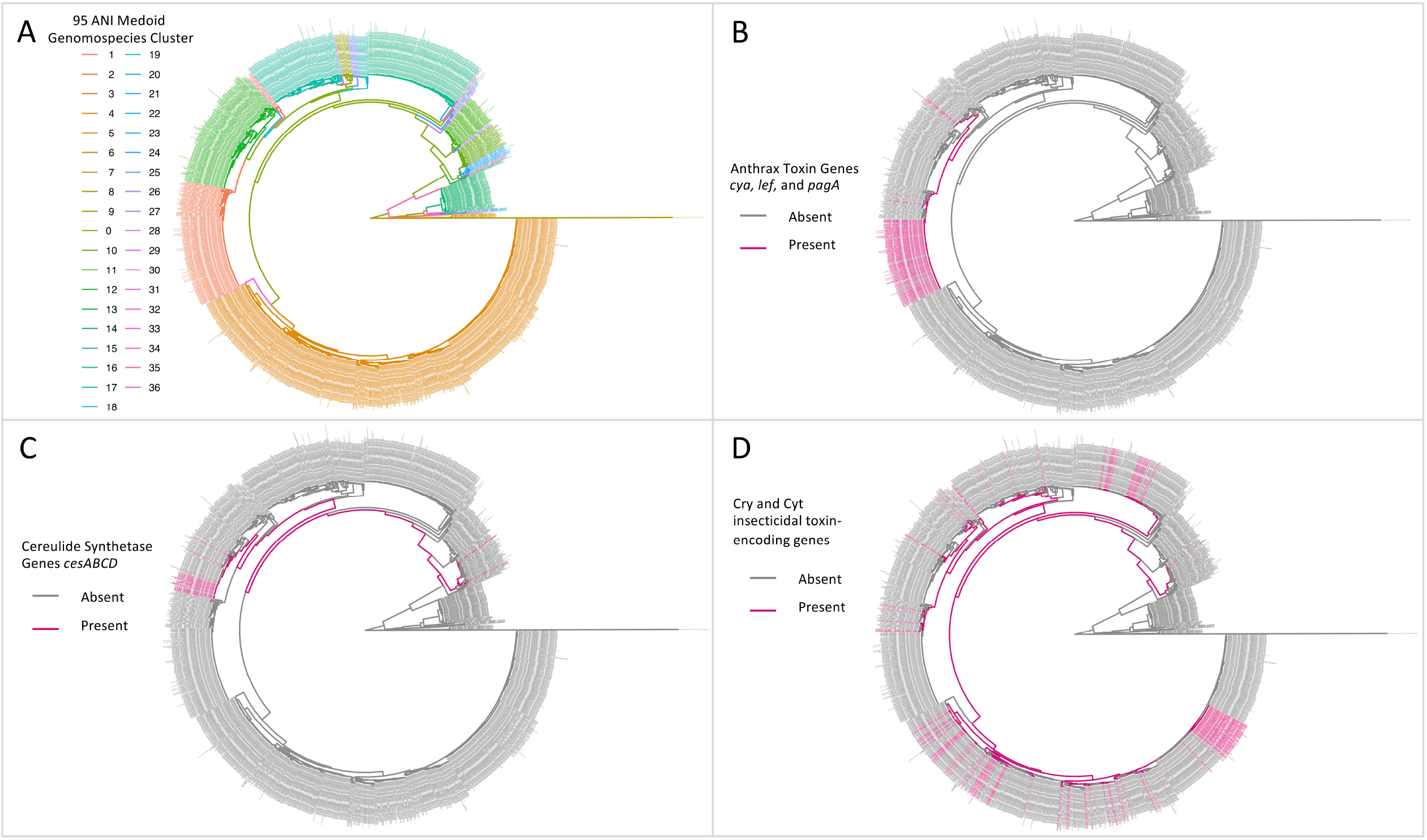
Maximum likelihood phylogenies of 2,218 *B. cereus* group genomes with N50 > 20 Kbp. Tip and branch labels are colored by (A) genomospecies assignment using medoid genomes of genomospecies clusters formed at the widely used genomospecies threshold of 95 ANI (clusters are arbitrarily numbered), and presence (pink) and absence (gray) of (B) anthrax toxin genes *cya*, *lef*, and *pagA*, (C) cereulide synthetase encoding *cesABCD*, and (D) one or more previously described Cry or Cyt insecticidal toxin-encoding genes. Phylogenies were constructed using core SNPs identified in 79 single-copy orthologous gene clusters present in 2,231 *B. cereus* group genomes. The type strain of “*B. manliponensis*” (i.e., the most distantly related member of the group) was treated as an outgroup on which each phylogeny was rooted. Virulence genes (*cya*, *lef*, and *pagA*; *cesABCD*) were detected using BTyper version 2.3.2 (default thresholds), while insecticidal toxin-encoding genes were detected using BtToxin_scanner version 1.0 (default settings; presence and absence of high-confidence, previously known Cry- and Cyt-encoding genes are shown, with predicted putative novel insecticidal toxin-encoding genes excluded).

Additionally, genes required for the production of anthrax toxin have been described in not only *B. anthracis*, but in isolates which share phenotypic characteristics often associated with “*B. cereus*” (e.g., motility, gamma bacteriophage resistance) as well.^10, 24, 37–40^ Despite the common assertion *that B. anthracis* is a clonal species with low diversity,^71–73^ the species cluster formed by *B. anthracis* at 95 ANI encompasses several lineages which fall outside of the highly similar one most commonly associated with anthrax illness (Figures 3A and 3B). Furthermore, even at the widely accepted genomospecies threshold of 95 ANI, nearly all (145 of 149; 97.3%) genomes possessing the anthrax toxin encoding genes (i.e., *cya*, *lef*, and *pagA*) were found to belong to the *B. anthracis* reference genome genomospecies cluster, including three of the seven genomes submitted to NCBI’s RefSeq database as anthrax-causing “*B. cereus*” (Figures 3A and 3B and Supplementary Table S6). These three genomes most closely resembled the *B. anthracis* reference genome, but also shared ≥95 ANI with the *B. paranthracis* type strain genome (Supplementary Table S6). The remaining four genomes derived from other anthrax-causing “*B. cereus*” strains most closely resembled the *B. tropicus* type strain, shared ≥95 ANI with the *B. paranthracis* type strain, and shared between 94 and 95 ANI with the *B. anthracis* species reference genome (Supplementary Table S6). This separation of anthrax-causing *B. cereus* group genomes into two genomospecies clusters at 95 ANI was maintained when medoid genomes were used in lieu of type/reference genomes as well (Figure 2A2 and Supplementary Table S6). As such, several anthrax-causing “*B. cereus*” strains are technically still *B. anthracis*, even by the current genomospecies definitions (i.e., ≥95 ANI relative to the *B. anthracis* species reference genome; Figures 3A and 3B) and despite having a mosaic of phenotypic characteristics attributed to *B. cereus s.s.* and *B. anthracis*.

Heterologous presentation within the genomospecies or lineage with which it is associated, as well as presence in additional genomospecies, is not reserved for anthrax toxin production. Emetic “*B. cereus*” has been designated as such by its ability to produce cereulide, a highly heat- and pH-resistant toxin responsible for a foodborne illness characterized by symptoms of vomiting.^1, 22, 74^ At a genomospecies threshold of ≥95 ANI, all 30 emetic “*B. cereus*” genomes most closely resembled the *B. paranthracis* type strain. All emetic “*B. cereus*” genomes were confined to a single genomospecies cluster when medoid genomes were used, and were interspersed among genomes which lacked *cesABCD* and are hence likely incapable of producing emetic toxin (Figures 3A and 3C and Supplementary Table S1). *cesABCD* were detected in five genomes representing two additional medoid-based genomospecies clusters at ≥95 ANI as well (Figures 3A and 3C and Supplementary Table S1). One of these genomospecies clusters contained the type strains of *B. weihenstephanensis* and *B. mycoides*, which is unsurprising considering cereulide-producing *B. weihenstephanensis* has been isolated in rare cases.^75, 76^ However, two genomes categorized previously as emetic “*B. weihenstephanensis*” belonged to a completely separate genomospecies cluster at a 95 ANI threshold (Figures 3A and 3C and Supplementary Table S1).

The Cry and Cyt insecticidal proteins associated with popular biocontrol agent *B. thuringiensis* (i.e., Bt toxins), which can be plasmid-mediated, are plagued by similar issues, as *B. thuringiensis* has historically been differentiated from *B. cereus s.s.* by its ability to produce insecticidal toxins (e.g., Cry and Cyt toxins).^77^ However, genes known to encode these insecticidal toxins were detected in nine of the 21 *B. cereus* group type strain genomospecies clusters at the widely used genomospecies threshold of 95 ANI (*B. albus*, *B. anthracis*, *B. cereus s.s., B. mycoides, B. paranthracis, B. thuringiensis, B. toyonensis, B. tropicus,* and *B. wiedmannii*; Figures 3A and 3D). These results are consistent with previous findings, as Bt toxin production has been previously attributed to numerous *B. cereus* group lineages.^69, 77, 78^

### ANI-based comparisons to medoid genomes using a lowered genomospecies threshold of ≈92.5 eliminate the species overlap problem for *B. anthracis* and its neighboring lineages

Numerous bacterial lineages have showcased a breakpoint in core genome similarity which is close to a threshold 95 ANI. As such, the 95 ANI cutoff has been proposed to serve as an adequate metric of delineation for many bacterial species.^15^ However, pairwise ANI values for a significant proportion of *B. cereus* group genomes, particularly *B. anthracis* and neighboring lineages, fall within the 93-95 ANI range, with a breakpoint in core genome similarity occurring close to 92.5 ANI (Figure 4 and Supplementary Figure S2). These characteristics, including lack of a natural breakpoint in core genome similarity at 95 ANI and a breakpoint at ≈92.5 ANI, were maintained when genomes were removed at quality and contamination filtering thresholds of varying stringency (Supplementary Figure S1). Using a 92.5 ANI breakpoint for genomospecies assignment, rather than 95, nearly eliminates the species overlap problem: only six of 2,231 genomes (0.269%) were assigned to 2 or more medoid-based genomospecies clusters at a hard threshold of 92.5 ANI (Figure 2 B2 and Supplementary Table S7). This can be compared to 18.2% and 66.2% of genomes assigned to multiple genomospecies clusters at 95 ANI when medoid genomes and species type strain/reference genomes were used, respectively (Figure 2 and Supplementary Tables S3 and S4).

**Figure 4.**
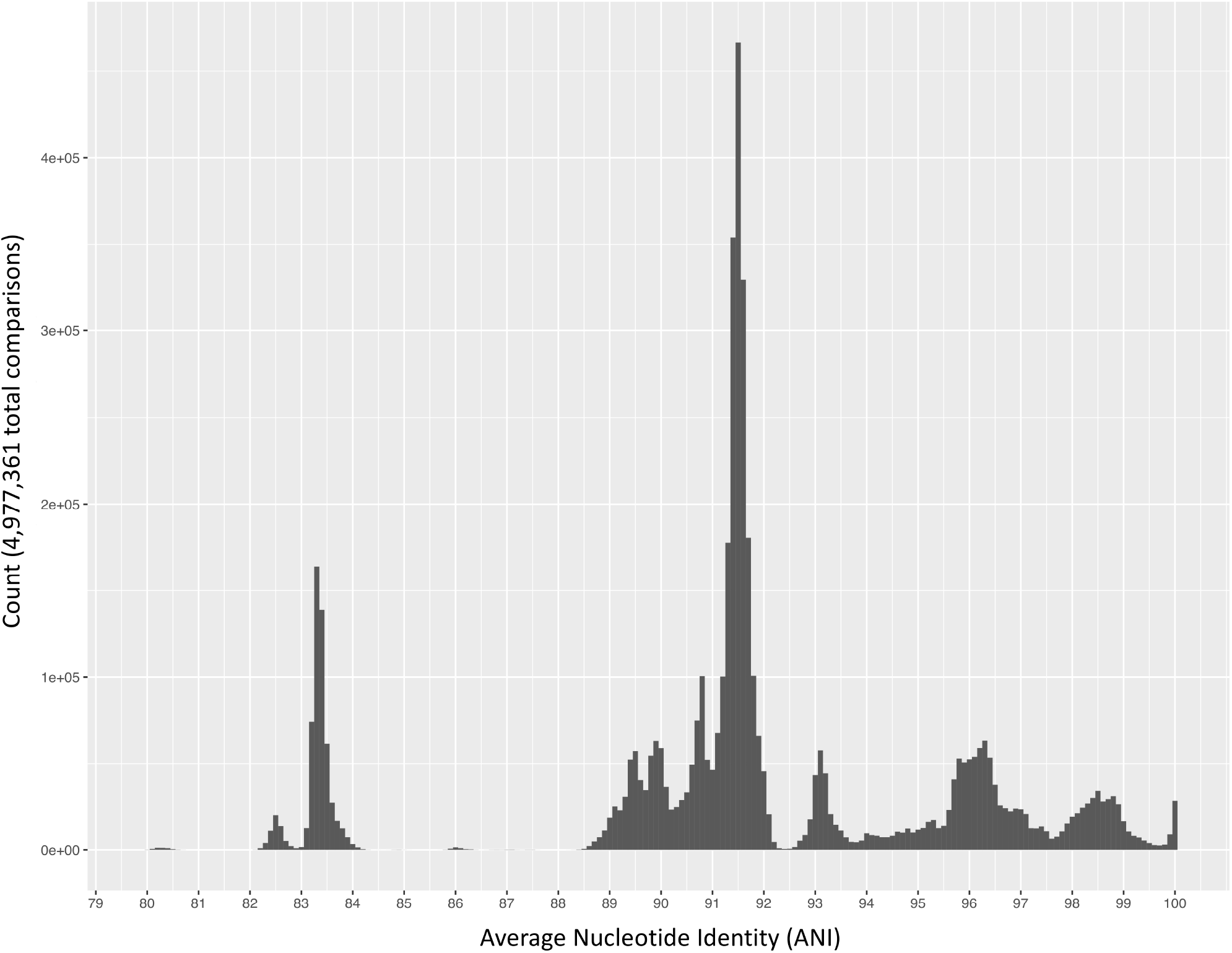
Histogram of pairwise average nucleotide identity (ANI) values calculated between 2,231 *B. cereus* group genomes downloaded from NCBI’s RefSeq database. FastANI version 1.0 was used to calculate all pairwise ANI values. For histograms of pairwise ANI values calculated between genomes meeting additional quality thresholds, or colored according to closest species type strain/reference genome at a traditional ≥95 ANI threshold, see Supplementary Figures S1 and S2, respectively.

Additionally, at 92.5 ANI, the total number of *B. cereus* group genomospecies clusters is reduced to 18, compared to 36 genomospecies clusters formed using medoid genomes at 95 ANI (Figures 3A and 5 and Supplementary Tables S4 and S7). At a threshold of 92.5 ANI, all genomes in which anthrax toxin-encoding genes were detected were confined to a single genomospecies cluster (referred to here as *B. mosaicus*; see Discussion section). Cereulide synthetase genes *cesABCD* were confined to two genomospecies clusters (*B. mosaicus* and *B. mycoides*; see Discussion section), while known Cry and Cyt genes commonly associated with *B. thuringiensis* were detected in four genomospecies clusters (*B. cereus s.s.*, *B. mosaicus*, *B. mycoides*, and *B. toyonensis*; see Discussion section).

**Figure 5.**
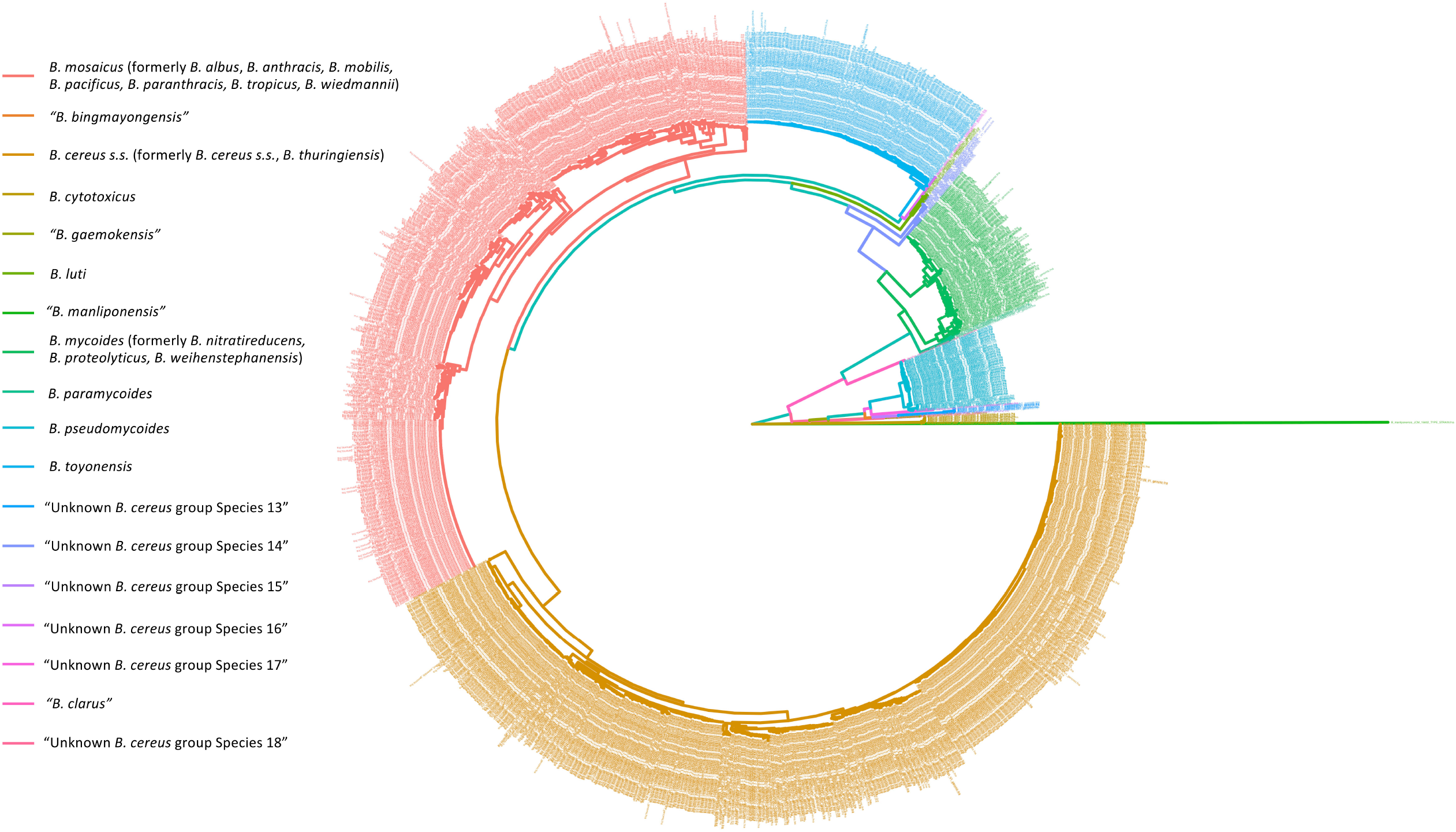
Maximum likelihood phylogeny of 2,218 *B. cereus* group genomes with N50 > 20 Kb. Tip and branch labels are colored by genomospecies assignment using medoid genomes of genomospecies clusters formed at proposed genomospecies threshold 92.5 ANI. Phylogeny was constructed using core SNPs identified in 79 single-copy orthologous gene clusters present in 2,231 *B. cereus* group genomes. The type strain of “*B. manliponensis*” (i.e., the most distantly related member of the group) was treated as an outgroup on which the phylogeny was rooted.

Notably, even at a lowered threshold of 92.5 ANI, seven genomospecies clusters did not possess type strains or reference genomes of any published species (Supplementary Table S7), indicating that putative novel *B. cereus* group genomospecies may be present among publicly available genomes. One of these genomospecies clusters has recently been proposed as novel *B. cereus* group species “*B. clarus*”.^79^ The remaining six genomospecies, which are composed of environmental *B. cereus* group strains from soil and agricultural environments, have not previously been proposed as novel species (Supplementary Table S8).

## DISCUSSION

When applied to bacteria, the taxonomic concept of “species” is notoriously ambiguous, particularly in cases where taxonomy is intertwined with a mobilizable (e.g., plasmid-encoded) phenotype, and even more so when that phenotype is a well-established component of the medical or industrial lexicon. Taxonomic definitions based solely on phenotype lack nuance in the omics era, as they ignore potential underpinning genomic diversity which could be leveraged to provide a higher-resolution assessment of an isolate’s pathogenic potential or industrial utility. A notable example outside of the *B. cereus* group can be seen in botulinum neurotoxin [BoNT]-producing bacterial species, to which the *Clostridium botulinum* species label has historically been applied, despite the fact that multiple genomospecies are capable of BoNT production.^80^ Furthermore, taxonomy based on phenotype can be ambiguous—and even misleading—when a trait is lost, gained, or not widespread throughout a lineage. For example, it is currently unclear if emetic “*B. cereus*” can still be labeled as such if it loses plasmid-encoded genes responsible for cereulide production. Additionally, emetic symptoms are not exclusive to cereulide intoxication.^81^ The development of a taxonomic nomenclature just for the sake of taxonomic rigor, however, can be equally problematic when a particular bacterial lineage has deep roots in medicine or industry. For example, some *Escherichia coli* lineages and *Shigella* spp., which do not represent distinct genera at a genomic level,^15, 82^ may be identified and treated differently in a clinical setting.^82^ As such, their current taxonomic designations are readily interpretable and actionable in the medical and public health communities, despite genomic inconsistencies reflected in their nomenclature.

An ideal taxonomic nomenclature for the *B. cereus* group should be easily interpretable by clinicians and public health officials, without sacrificing the resolution provided by WGS and other contemporary technologies. Several previous publications describing *B. cereus* group members which exhibit genotype-phenotype incongruencies have appended the term “biovar” to species names to denote phenotypes of interest (e.g., anthrax-causing “*B. cereus*” as *B. cereus* biovar anthracis; Cry-producing *B. wiedmannii* as *B. wiedmannii* biovar thuringiensis).^38, 78^ A taxonomic nomenclature for the *B. cereus* group is thus proposed, consisting of the following components: (i) an amended collection of genomospecies names, corresponding to the resolvable genomospecies clusters obtained at the *B. cereus* group core genome breakpoint of ≈92.5 ANI shown here; (ii) a formal collection of subspecies names, which can be used to account for well-established lineages of medical importance; and (iii) a formalized and extended collection of biovar terms, which can account for phenotype heterogeneity (Figure 6; note that a recently proposed “genomovar” framework for *B. cereus s.s.* and genomes classified as *B. thuringiensis* using the species type strain genome^83^ is not adopted here, due to the lack of a genomospecies boundary for *B. cereus s.s.* and *B. thuringiensis* [shown here and elsewhere,^17–19^ including the paper proposing the genomovar framework^83^], as well as the lack of a standardized species definition for *B. thuringiensis* [i.e., some studies have defined *B. thuringiensis* as any *B. cereus* group species capable of producing insecticidal toxins,^69^ while others have defined it based on similarity to the species type strain genome^83^]).

**Figure 6.**
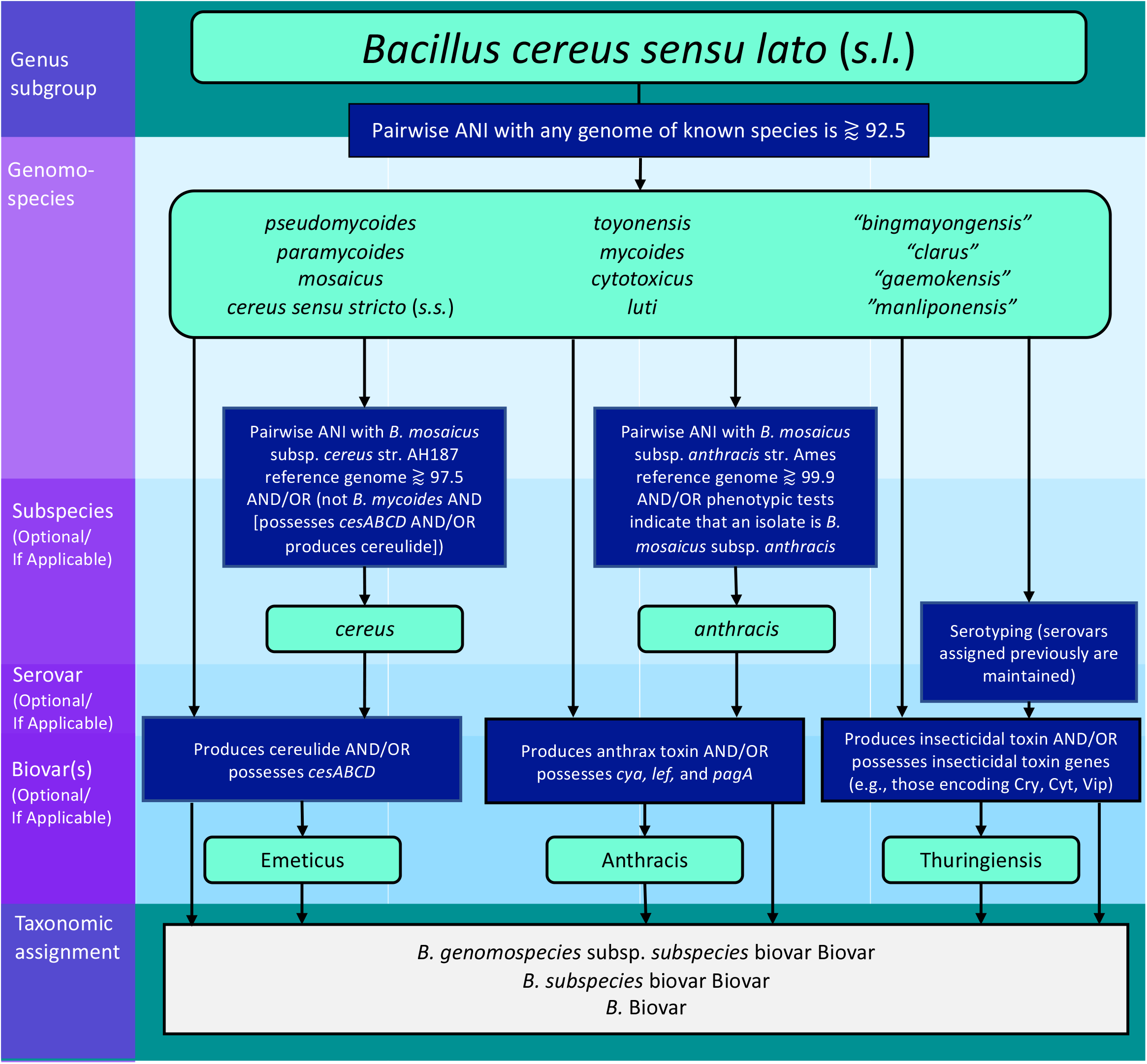
Taxonomic hierarchy for the proposed *B. cereus* group nomenclature. Taxonomic levels are listed in the left margin, with levels which are optional/not applicable to all organisms denoted as such. Rounded boxes shaded in light green correspond to possible taxonomic designations at their respective level, while blue boxes correspond to requirements an isolate and/or its genome must meet to be assigned that designation. Possible forms which the final taxonomic assignment can take can be found in the gray box at the bottom of the chart.

### A formal proposal of a novel taxonomic nomenclature for the *B. cereus* group

#### A. Genomospecies

The *B. cereus* group currently consists of eight genomospecies clusters (denoted I – VIII) which encompass published *B. cereus* group species, four genomospecies (denoted ix – xii) which encompass putative *B. cereus* group species that have already been proposed in the literature, and six genomospecies (denoted xiii – xviii) which may represent unknown putative genomospecies that have yet to be proposed (Figure 5). A genome belongs to a genomospecies if it shares ≧ 92.5 ANI with the genomospecies medoid genome (Supplementary Table S7). Due to the resolvability of genomospecies clusters at this threshold, it follows that (i) a genome does not belong to a genomospecies if it shares ≦ 92.5 with the genomospecies medoid genome; (ii) two genomes belong to the same genomospecies if they share ≧ 92.5 ANI with each other; (iii) two genomes belong to different genomospecies if they share ≦ 92.5 ANI with each other (i.e., in practice, one does not need to use a genomospecies medoid genome for genomospecies assignment, but rather any genome of known genomospecies; see Supplementary Tables S1 [genomospecies assignments for all publicly available *B. cereus* group genomes] and S7 [genomospecies assignments of *B. cereus* group type strain genomes]). When written, genomospecies names immediately follow the genus name (*Bacillus* or *B.*) and are italicized and lower-case.

### Published genomospecies

#### I. Bacillus pseudomycoides

The *B. pseudomycoides* genomospecies cluster contained 111 genomes, including the genome of species type strain *B. pseudomycoides* str. DSM 12442. All genomes previously classified as *B. pseudomycoides* relative to the type strain at a threshold of 95 ANI remain in this genomospecies, and no additional genomes belong to the genomospecies. As such, this genomospecies remains consistent with its previous classification, and its previous name remains unchanged.

#### II. Bacillus paramycoides

The *B. paramycoides* genomospecies cluster contained six genomes, including the genome of species type strain *B. paramycoides* str. NH24A2. All genomes previously classified as *B. paramycoides* relative to the type strain at a threshold of 95 ANI remain in this genomospecies, and no additional genomes belong to the genomospecies. As such, this genomospecies remains consistent with its previous classification, and its previous name remains unchanged.

#### III. Bacillus mosaicus

The *B. mosaicus* genomospecies contained 722 genomes, including type strains and reference genomes of species formerly known as *B. albus* (now *B. mosaicus* str. N35-10-2), *B. anthracis* (now *B. mosaicus* subsp. *anthracis* str. Ames; see sections “Subspecies” and “Biovars” below)*, B. mobilis* (now *B. mosaicus* str. 0711P9-1), *B. pacificus* (now *B. mosaicus* EB422), *B. paranthracis* (now *B. mosaicus* str. MN5)*, B. tropicus* (now *B. mosaicus* N24), and *B. wiedmannii* (now *B. mosaicus* FSL W8-0169). Additionally, all members of the lineage formerly known as emetic “*B. cereus*” belong to *B. mosaicus* (see sections “Subspecies” and “Biovars” below). While the species formerly known as *B*. *anthracis* is the oldest described former species in this group, it is not proposed as the genomospecies name, as doing so could lead to incorrect assumptions of an isolate’s anthrax-causing potential. As such, the proposed genomospecies name (*mosaicus*) is chosen to reflect the diversity of lineages and phenotypes present among members of this genomospecies. All genomes previously assigned to the aforementioned former species using their respective type strain or reference genomes at a threshold of 95 ANI belong to *B. mosaicus*.

#### IV. Bacillus cereus sensu stricto (s.s.)

The *B. cereus s.s.* genomospecies contained 949 genomes, including those of type strains *B. cereus s.s.* (*B. cereus s.s.* str. ATCC 14579) and former species *B. thuringiensis* (now *B. cereus s.s.* serovar berliner biovar Thuringiensis str. ATCC 10792; see section “Biovars” below). *B. cereus s.s.* was chosen as the genomospecies name, with Thuringiensis proposed as a biovar to account for phenotypic heterogeneity within *B. cereus s.s.*, as well as the presence of insecticidal toxins in other genomospecies (see section “Biovars” below). All genomes previously assigned to the species *B. cereus s.s.* and former species *B. thuringiensis* at a 95 ANI threshold using these type strains belong to the *B. cereus s.s.* genomospecies.

#### V. Bacillus toyonensis

The *B. toyonensis* genomospecies contained 230 genomes, including the type strain of *B. toyonensis* (*B. toyonensis* str. BCT-7112). All genomes previously classified as *B. toyonensis* relative to the type strain at a threshold of 95 ANI remain in this genomospecies, and no additional genomes belong to the genomospecies. As such, this genomospecies remains consistent with its previous classification, and its previous name remains unchanged.

#### VI. Bacillus mycoides

The *B. mycoides* genomospecies contained 164 genomes, including the type strain of *B. mycoides* (*B. mycoides* str. DSM 2048), former species *B. nitratireducens* (now *B. mycoides* str. 4049), former species *B. proteolyticus* (now *B. mycoides* str. TD42), and former species *B. weihenstephanensis* (now *B. mycoides* str. WSBC 10204). Additionally, all members of the lineages formerly known as emetic *B. weihenstephanensis* belong to *B. mycoides* (see section “Biovars” below). *B. mycoides* was selected as the genomospecies name, as it is the oldest of published former species described in this cluster (and remains consistent with taxonomic changes recently proposed by others).^68^ All genomes previously assigned to the aforementioned species using their respective type strain or reference genomes and a threshold of 95 ANI belong to *B. mycoides*.

#### VII. Bacillus cytotoxicus

The *B. cytotoxicus* genomospecies contained 14 genomes, including the type strain of *B. cytotoxicus* (*B. cytotoxicus* str. NVH 391-98). All genomes previously classified as *B. cytotoxicus* relative to the type strain at a threshold of 95 ANI remain in this genomospecies, and no additional genomes belong to the genomospecies. As such, this genomospecies remains consistent with its previous classification, and its previous name remains unchanged.

#### VIII. Bacillus luti

The *B. luti* genomospecies contains nine genomes, including the type strain of *B. luti* (*B. luti* str. TD41). All genomes previously classified as *B. luti* relative to the type strain at a threshold of 95 ANI remain in this genomospecies, and no additional genomes belong to the genomospecies. As such, this genomospecies remains consistent with its previous classification, and its previous name remains unchanged.

### Previously proposed putative species

The following four putative *B. cereus* group genomospecies which have been proposed previously remain unchanged:

**ix. *“B. bingmayongensis”*** (including type strain *“B. bingmayongensis”* str. FJAT-13831)

**x. “*B. gaemokensis”*** (including type strain *“B. gaemokensis”* str. KCTC 13318)

**xi. “*B. manliponensis”*** (including type strain *“B. manliponensis”* str. JCM 15802)

**xii. “*B. clarus”*** (including type strain “*B. clarus”* str. ATCC 21929)

### Putative novel species

The six putative genomospecies clusters xiii-xviii listed in Supplementary Table S8 have not been proposed as novel species in the literature, and thus may eventually be adopted as novel species following rigorous genotypic and phenotypic characterization and peer-reviewed publication. Future proposed novel *B. cereus* group species should (i) share < 92.5 ANI with all *B. cereus* group genomes, and (ii) share ≥ 97% 16S rDNA similarity with known *B. cereus* group species (a definition used in previous studies).^19^

#### B. Subspecies

We propose the adoption of the following two subspecies terms to ensure that the medically important lineages formerly known as *B. anthracis* and emetic “*B. cereus*” are still interpretable outside of a strictly taxonomic context. When written, subspecies names are italicized, lower-case, and can optionally (i) be appended to the species name, after the non-italicized delimiter “subspecies” or “subsp.”, prior to a serotype designation (if applicable); or (ii) follow the genus name (*Bacillus* or *B.*) directly, with the species name omitted, prior to a serotype designation (if applicable).

a. *B. mosaicus* subspecies *anthracis* (can also be written as *B. mosaicus* subsp. *anthracis*; *B. anthracis*): refers to the comparatively clonal lineage of former species B. *anthracis* commonly associated with anthrax illness. Isolates which are assigned to this subspecies should (i) exhibit distinguishing phenotypic characteristics (e.g., lack of motility, lack of hemolysis on Sheep RBC agar) associated with the classical definition of *B. anthracis* as outlined in the Bacteriological Analytical Manual (BAM) chapter on *B. cereus*,^10^ and/or (ii) share ≧ 99.9 ANI with former species reference genome *B. anthracis* str. Ames (now *B. mosaicus* subsp. *anthracis*; NCBI RefSeq Accession GCF_000007845.1), as Jain, et al.^15^ identified this as a threshold for this closely related lineage, a result which we replicated here. The use of the term “subspecies *anthracis*” does not indicate whether an isolate produces anthrax toxin or possesses the machinery required for the synthesis of anthrax toxin or not (see “biovar Anthracis” below for further clarification).
b. *B. mosaicus* subspecies *cereus* (can also be written as *B. mosaicus* subsp. *cereus*; *B. cereus*): refers to the lineage formerly known as emetic “*B. cereus*”. All genomes possessing cereulide synthetase genes (*cesABCD*) which did not belong to the *B. mycoides* species cluster (see “Species” section above) shared ≥97.5 ANI with the emetic reference strain formerly known as *B. cereus* str. AH187 (now *B. mosaicus* subsp. *cereus* biovar Emeticus; NCBI RefSeq Accession GCF_000021225.1). As such, isolates which are assigned to this subspecies (i) produce cereulide and belong to the species *B. mosaicus*, (ii) possess the cereulide synthetase biosynthetic gene cluster and belong to the species *B. mosaicus*, and/or (iii) share ≧97.5 ANI with emetic reference genome *B. cereus* str. AH187 (now *B. mosaicus* subsp. *cereus* biovar Emeticus; NCBI RefSeq Accession GCF_000021225.1). The use of the term “subspecies *cereus*” does not indicate whether an isolate produces cereulide or possesses the machinery required for the synthesis of cereulide or not (see “Biovar Emeticus” below for further clarification).

#### C. Biovars

To account for phenotypes of clinical and industrial importance which can be distributed across species and heterogeneous in their appearance in individual lineages, we propose the biovars listed below. While phenotypic evidence of a trait assigned to a biovar is ideal, biovars can also be predicted at the genomic level. When written, (i) the first letter of the biovar is capitalized; (ii) the biovar name is not italicized; (iii) the biovar is appended to the end of a species, subspecies (if applicable), or serotype name (if applicable), following the non-italicized delimiter “biovar”; (iv) if applicable, multiple biovars follow the non-italicized, plural delimiter “biovars”, are listed in alphabetical order, and are each separated by a comma and a single space; (v) biovar(s) may follow the genus name (*Bacillus* or *B.*) directly, with the species, subspecies (if applicable), and serotype (if applicable) names omitted.

a. **biovar Anthracis**: can be applied to an isolate (i) known to produce anthrax toxin (preferred), and/or (ii) possess anthrax toxin encoding genes *cya*, *lef,* and *pagA*. Capsular genes (e.g., *cap*, *has*, *bps*)^30, 31, 84^ are deliberately excluded from the definition of the Anthracis biovar as a conservative measure. This is to avoid cases in which an isolate might possess anthrax toxin genes but no known capsule synthesis genes, despite the ability to synthesize a capsule via novel capsule synthesis mechanisms. Published examples of this biovar are: *B. mosaicus* subsp. *anthracis* biovar Anthracis (i.e., anthrax-causing members of the classical “clonal” lineage often associated with anthrax disease; can also be written as *B. anthracis* biovar Anthracis or *B.* Anthracis); *B. mosaicus* biovar Anthracis (i.e., anthrax-causing lineages formerly known as “anthrax-causing *B. cereus*”; can also be written as *B.* Anthracis).
b. **biovar Emeticus**: can be applied to an isolate known to produce cereulide (preferred) and/or possess genes encoding cereulide synthetase (*cesABCD*). Published examples of this biovar are: *B. mosaicus* subsp. *cereus* biovar Emeticus (i.e., cereulide-producing lineages formerly known as emetic “*B. cereus*”; can also be written as *B. cereus* biovar Emeticus or *B.* Emeticus); *B. mycoides* biovar Emeticus (i.e., cereulide-producing lineages formerly known as “emetic *B. weihenstephanensis*”; can also be written as *B.* Emeticus).
c. **biovar Thuringiensis**: can be applied to an isolate known to produce one or more insecticidal/Bt toxins (preferred) and/or possess genes known to encode insecticidal toxins (e.g., genes encoding Cry, Cyt, or Vip toxins). Examples of this biovar include *B. mosaicus* biovar Thuringiensis and *B. cereus s.s.* biovar Thuringiensis (both of which can be written as *B.* Thuringiensis).

## CONCLUSION

The nomenclature proposed here offers numerous advantages over previous taxonomic conventions. Most importantly, it is consistent; it provides an explicit, standardized framework for naming and classifying members of the *B. cereus* group using genomic and/or phenotypic methods, and it resolves previous ambiguities in the scientific community (e.g., whether *B. cereus* group isolates outside of the “clonal” lineage often associated with anthrax disease but still within commonly employed genomospecies thresholds, including those capable of causing anthrax,^40, 85, 86^ are *B. anthracis*; whether “*B. thuringiensis*” refers to any *B. cereus* group member which carries genes encoding insecticidal toxins,^69^ or to members of the *B. cereus* group which most closely resemble the *B. thuringiensis* species type strain;^83^ whether emetic “*B. cereus*” should be referred to as such, or as “*B. paranthracis*”^23^). Furthermore, genomes can now only be assigned to a single genomospecies (i.e., the species-overlap problem is eliminated), and a single, genomically informed ANI threshold for the proposal of novel genomospecies is proposed.

A second advantage of the proposed taxonomy is its backwards-compatibility with previous medical and industrial definitions of *B. cereus* group species; for example, with the taxonomy proposed here, any *B. cereus* group isolate capable of producing insecticidal crystal proteins can be referred to as *B.* Thuringiensis, which is in line with the traditional definition of the species.^69^ Specific lineages, however, can be accounted for through the incorporation of species and/or serovar names (e.g., *B. cereus s.s.* serovar berliner biovar Thuringiensis).

Additionally all isolates capable of producing anthrax toxin can be referred to as *B.* Anthracis, while members of the “clonal” anthrax lineage can continue to be referred to as *B. anthracis* (the subspecies short-form of *B. mosaicus* subsp. *anthracis*). We anticipate that these minor nomenclatural changes will remain interpretable and actionable to those in medicine, public health, and industry, while still remaining true to genomic definitions of bacterial species.

Finally, the proposed taxonomy is advantageous for its flexibility. The framework proposed here can be easily expanded to account for additional important lineages or phenotypes through the adoption of novel subspecies or biovars, respectively. For example, future biovars (i.e., biovar Cereus) can be proposed to describe *B. cereus* group members capable of causing diarrheal foodborne disease, as this form of disease involves multiple toxins and is not yet fully understood.^87^ The biovar nomenclature proposed here, along with revised genomospecies definitions and the proposal of novel subspecies, provides a standardized framework for *B. cereus* group classification, accounting for both phylogenomic diversity and phenotypic heterogeneity. An open-source, freely available command-line tool for characterizing *B. cereus* group genomes *in silico* using the framework proposed here can be found at: https://github.com/lmc297/BTyper3.

## ACKNOWLEDGMENTS

This material is based on work supported by the National Science Foundation Graduate Research Fellowship Program under grant no. DGE-1650441 and USDA National Institute of Food and Agriculture Hatch Appropriations under Project #PEN04646 and Accession #1015787.

## AUTHOR CONTRIBUTIONS

LMC performed all computational analyses. JK, LMC, and MW conceived the study and wrote the manuscript.

