## Supplementary Figure S1 for "Proposal of a taxonomic nomenclature for the *Bacillus cereus* group which reconciles genomic definitions of bacterial species with clinical and industrial phenotypes"

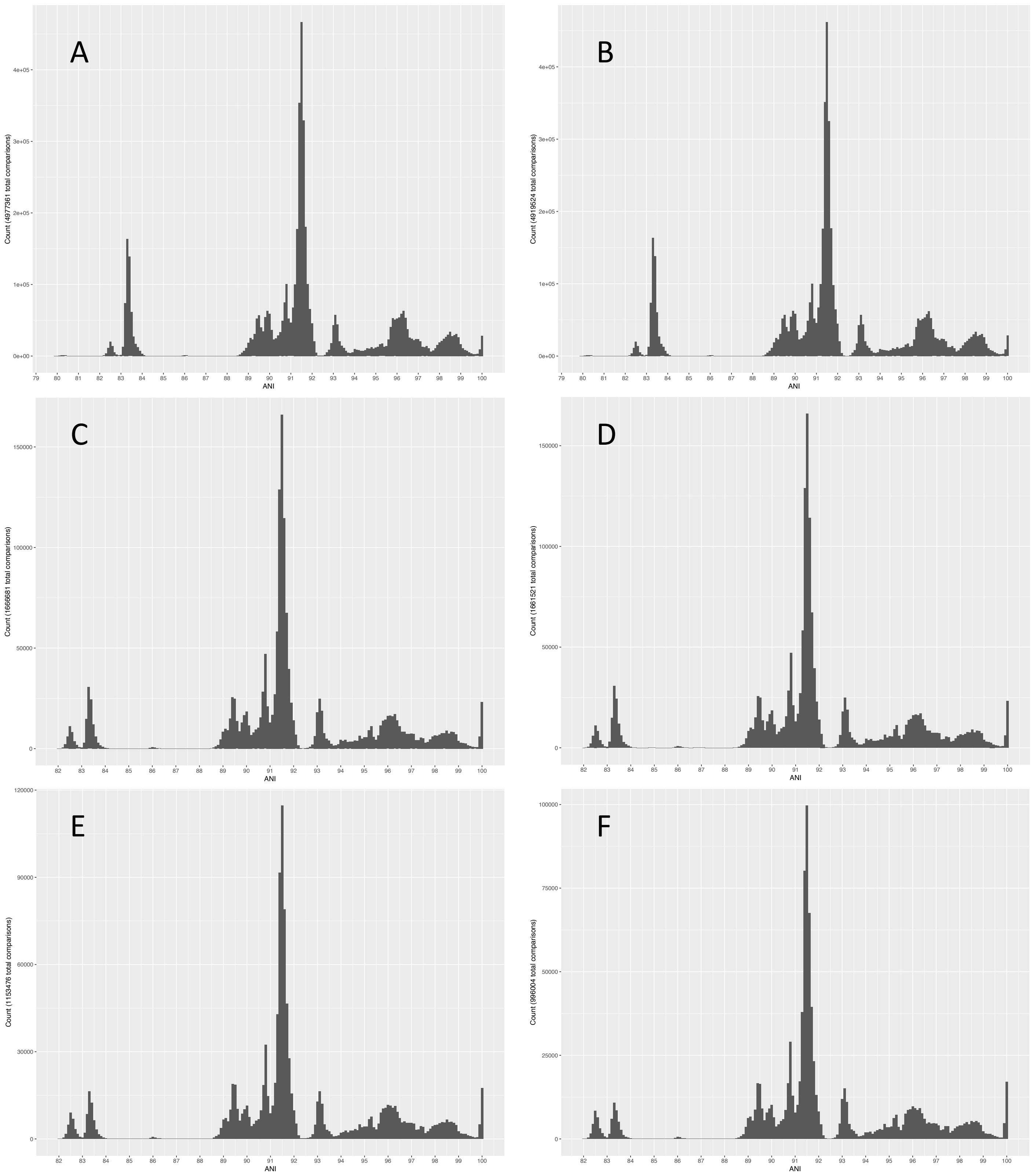


**Supplementary Figure S1.** Sample of histograms constructed using pairwise ANI values calculated between *B. cereus* group genomes meeting various quality thresholds. Because distribution breakpoints and shape were robust to the exclusion of genomes at all tested thresholds, the six histograms shown represent pairwise ANI values calculated between (A) raw genomes downloaded directly from NCBI's RefSeq database (i.e., the least stringent quality thresholds, with no filtering criteria employed; *n* = 2,231 genomes); (B) genomes with N50 > 20 Kb (the set of genomes from which medoid genomes were selected; *n* = 2,218); (C) genomes with N50 > 0 and no contigs assigned to a domain other than Bacteria (*n* = 1,291), (D) genomes with N50 > 20Kb and no contigs assigned to a domain other than Bacteria (*n* = 1,289), (E) genomes with N50 > 0 and no contigs assigned to a genus outside of *Bacillus* (*n* = 1,074), and (F) genomes with N50 > 100Kb and no contigs assigned to a genus outside of *Bacillus* (i.e., the most stringent quality thresholds tested; *n* = 998).
