## Supplementary Figure S2 for "Proposal of a taxonomic nomenclature for the *Bacillus cereus* group which reconciles genomic definitions of bacterial species with clinical and industrial phenotypes"

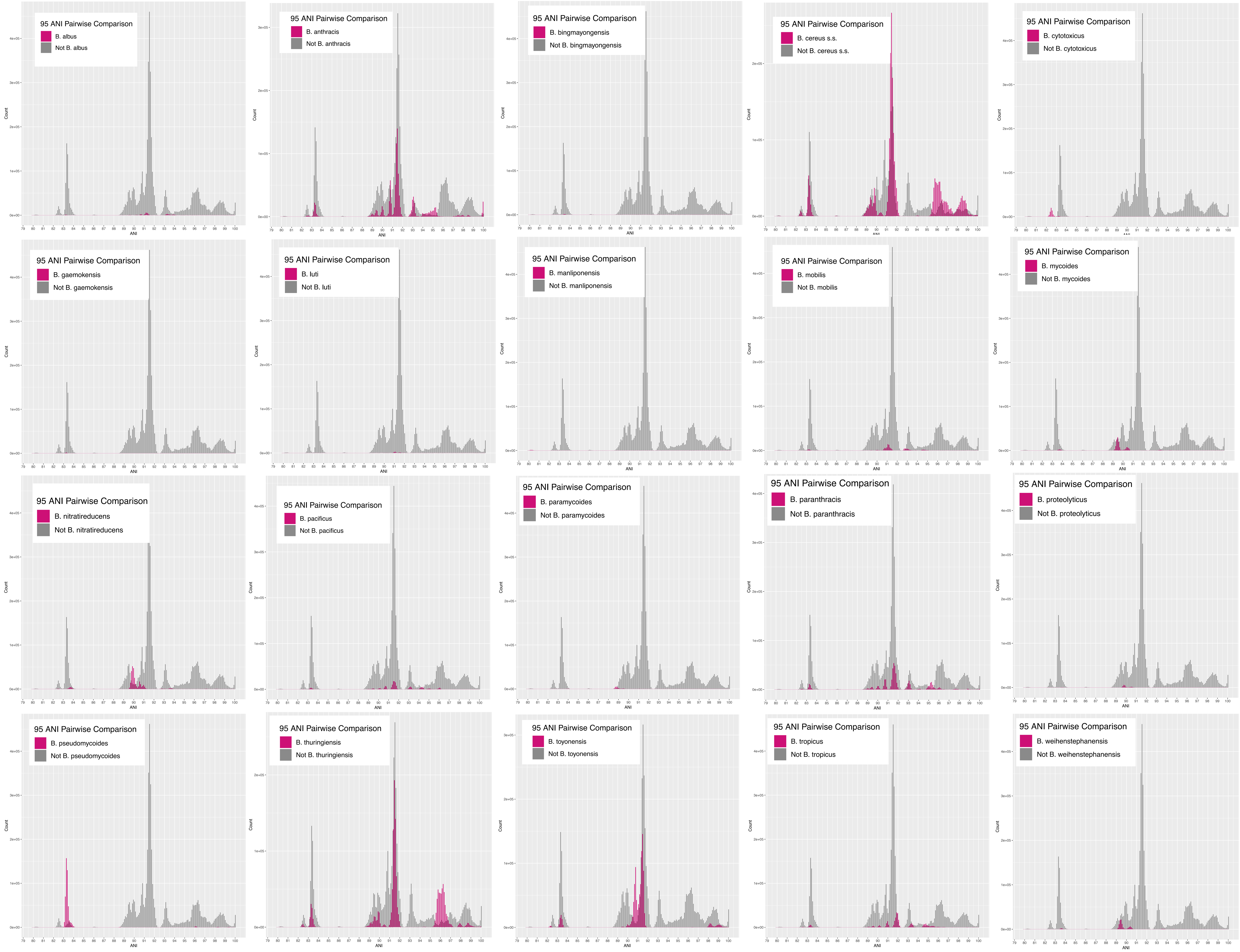


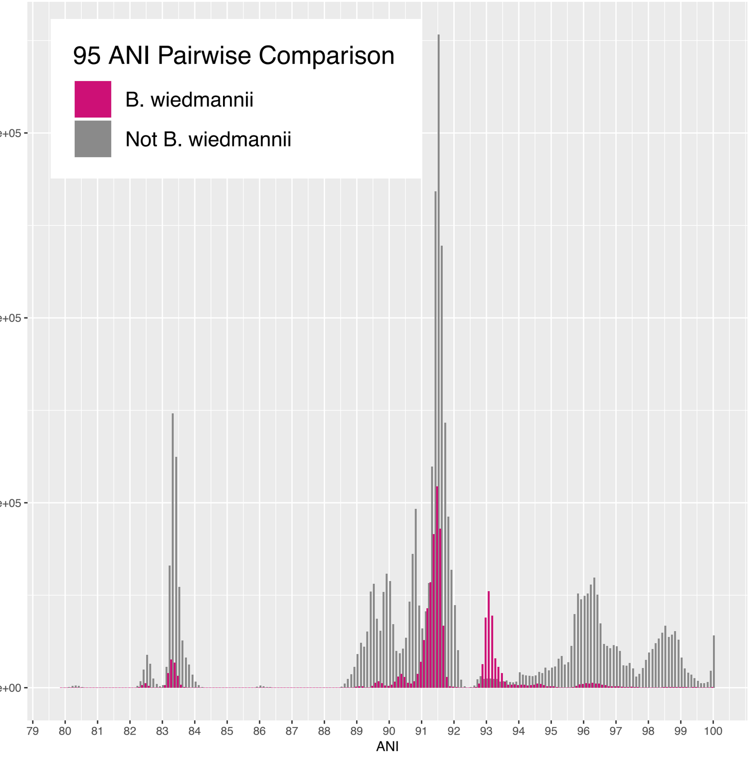


**Supplementary Figure S2.** Histograms of pairwise ANI values between 2,218 *B. cereus* group genomes with N50 > 20 Kbp. For each of 18 published species and 3 proposed putative species, ANI values calculated between two genomes for which one or more genome(s) shared ≥ 95 ANI with the species type strain or reference genome are colored in pink. ANI values calculated between two genomes for which both genomes shared < 95 ANI with the species type strain/reference genome are colored in gray. See Supplementary Table S2 for details regarding species type strain genomes.
